# Evolutionary history with chronic malnutrition enhances pathogen susceptibility at older ages

**DOI:** 10.1101/2024.10.04.616684

**Authors:** Saubhik Sarkar, Biswajit Shit, Joy Bose, Souvik De, Tadeusz J. Kawecki, Imroze Khan

## Abstract

Juvenile malnutrition is a global public health concern that negatively impacts the development and maturation of the immune system, leading to increased susceptibility to infectious diseases. Such adverse effects on immunity might increase with ageing, worsening disease conditions later in life. Furthermore, malnutrition may persist across generations, imposing strong natural selection to survive the nutrient shortage. However, it is unclear how the evolutionary history of ancestral generations with chronic malnutrition could influence pathogen resistance and infection susceptibility, as well as their age-specific changes in extant generations. To address this, we used *Drosophila melanogaster* populations adapted against chronic juvenile malnutrition and exposed them to a bacterial pathogen, *Providencia rettgeri*, during their early and late adulthood. Surprisingly, we observed that in populations adapted to chronic malnutrition, young flies survived infection better by tolerating the infection, while control flies displayed higher infection susceptibility despite carrying a similar pathogen load. However, this pattern in post-infection survival is reversed with ageing. There was no change in pathogen resistance, but evolved flies succumbed more to infection than control flies regardless of the input infection doses. Our study thus revealed new evolutionary insights into the development of contrasting early vs late-life immune strategies and age-specific vulnerabilities to infection as a function of early-life malnutrition.

## INTRODUCTION

Different ecological stressors often exacerbate each other’s effects on an individual’s health and Darwinian fitness. This is particularly the case for interactions between malnutrition, resulting from a diet of poor quality or insufficient quantity, and infectious disease (Wells et al., 2020). The increased susceptibility of nutritionally stressed individuals to infections may partly be caused by the impairment of immune function by food deprivation and/or lack of nutrients and altered nutrient sensing pathways during malnutrition (Bourke et al., 2016), exacerbating the disease conditions (Rytter et al., 2014). Such synergistically adverse effects may be particularly strong in juvenile animals. Because of their low competitiveness and the requirement for protein to grow, juveniles are often the first to face the adverse effects of malnourishment (Norris et al., 2022). At the same time, their immune system may not yet mature to counter particular pathogens for the first time. This is supported by direct experimental evidence from studies on species ranging from nematodes and fruit flies to murine models, where it has been shown that inadequate dietary intake at the juvenile stage reduces immune protection against invading pathogens and renders an individual more prone to pathogenic infection (Unckless et al., 2015; Chakravarty, 2022; Coutinho et al., 2008). In humans, too, malnourished children, soon after birth, might face a cycle of food scarcity, poor nutrition, and infections, proportionally increasing the risk of death once infected (Rice et al., 2000). Besides, they are also predisposed to common diseases such as diarrhoea, malaria, measles and chronic respiratory tract infections (Bryce et al., 2005), including SARS-COV-2 infections (Calder, 2021). Moreover, malnutrition during pre and post-natal development, including small body size at birth, can also lead to later-life morbidity and mortality, including reduced lifespan (Davis et al., 2016), altered metabolic function (Szostaczuk et al., 2020), and early-onset of age-related diseases (Barker, 1986; Hoffman et al., 2021). Since immune responses are inherently compromised with ageing, malnutrition at early ages in such cases might also act synergistically to exacerbate the severity of infection in older individuals (Khan et al., 2017; Shit et al. 2023), but the empirical evidence is limited.

Also, regular exposure to nutritional stress results in natural selection to favour mechanisms that alleviate its consequences (Kolss et al., 2009; Cavigliasso et al., 2020). Hence, the extent to which the evolutionary past with poor nutrition affects the outcome of pathogenic infections must be clarified. This is particularly relevant for many human populations grappling with inadequate nutrition for generations (Victora et al., 2008; Roseboom et al., 2006) and wild animals regularly getting exposed to suboptimal food (Birnie-Gauvin et al., 2017). They may evolve a better post-infection survival during early life as a general cross-resistance strategy against stressful environments (Le Bourg et al., 2008). Alternatively, they might experience a strong selection to overcome the fitness costs of such nutritional stress by evolving an optimal resource allocation to maintain or enhance traits with immediate fitness benefits at an early life. This may trade-off with somatic maintenance, immunity, and repair during future pathogen attacks at old ages, making them more vulnerable to diseases due to antagonistic pleiotrophy effects (Dasgupta et al., 2022). A previous experiment by Vijendraverma et al. (2015 with the fruit fly *Drosophila melanogaster* suggested that populations selected for increased survival to chronic malnutrition were more susceptible to food-borne infections caused by a gut pathogen *Pseudomonas entomophila*, and the gut was implicated as a focus of trade-off between nutrition acquisition and resistance against food-borne pathogens. However, it is unclear if the effect is consistent across pathogens and modes of infection. Also, it remains unexplored if adaptation to early-life malnutrition in these fly populations may lead to stronger trade-offs with post-infection survival later in life.

In this work, we tested these possibilities using *D. melanogaster* populations genetically adapted against chronic larval malnutrition for ∼275 generations of experimental evolution (Erkosar *et al*., 2023). These populations evolved significantly faster growth rates than their control populations under malnourishment (Kolss et al., 2009; Vijendravarma and Kawecki, 2013). We report that while these malnutrition-adapted populations showed an increased tolerance against systemic infection caused by a natural fly pathogen, *Providencia rettgeri* at a young age, they became more susceptible to bacterial infection when they were old, despite no changes in bacterial load. Our work thus unveiled the possibility of age-dependent antagonistic pleiotropy with infection outcomes during adaptation against poor nutrition.

## MATERIALS AND METHODS

### Fly populations

We used replicated *Drosophila melanogaster* populations adapted to chronic juvenile malnourishment (See Kolss et al., 2009 for details). The control populations have been maintained on standard food (15 g agar, 30 g sucrose, 60 g glucose, 12.5 g dry yeast, 50 g cornmeal, 0.5 g MgSO_4_, 0.5 g CaCl_2_, 30 mL ethanol, 6 mL propionic acid and 1 g nipagin per litre of water), whereas selected populations have been reared on a poor larval food regime, containing 25% of the amounts of sugars, yeast and cornmeal of the standard food, imposing severe nutritional stress at the pre-adult stage for over ∼275 generations. Although both regimes included six replicate populations derived from a single, laboratory-adapted base population, we only used three randomly chosen replicate populations from each regime for logistical reasons. Upon emergence, we transferred the adult flies to standard food. To generate standardised experimental flies, we used standard food to rear parental generations of control and selected flies for one generation, minimizing transgenerational effects, then collected experimental flies in the subsequent generation. Since deprivation of developmentally essential nutrients decreased significantly more the lifespan of female *Drosophila* than males (Wu et al., 2019), we performed our assays with females only. For each replicate population, we collected adult females and held them as virgins at a density of 15 flies/vial. We transferred the flies to fresh media vials every ∼2 days until they were assayed. Since *Drosophila* females undergo reproductive senescence within 30 days post-eclosion (Shit et al., 2022), hence we used 2-day vs 30-day-old flies (post-eclosion) as young and old flies, respectively.

### Bacterial infection and assays

We used a natural fly pathogen, the gram-negative bacteria *Providencia rettgeri* (*strain Dmel*, Hanson et al., 2019), to cause septic infection in our experimental flies, using a protocol described in Shit et al. (2022). At each age point, we pricked individual flies into their thoracic region using a 0.1mm minuten pin (Fine Science Tool) dipped in bacterial suspension adjusted to 4 different concentrations, namely 1, 0.5, 0.1 and 0.05 OD (measured at 600 nm, originally derived from a 10 ml of overnight-grown 1 OD culture of *P. rettgeri*). We also pricked flies with sterile phosphate-buffered saline (1X) as a procedural control (or sham infection). We infected a total of n=75 females/ infection dose/ age/ selection regime/ replicate populations, and redistributed them in food vials in a batch of 15 individual flies (we thus had five independent replicate vials for each infection dose, selection regime, replicate population and age points).

Following this, from each vial, we randomly removed three flies and measured their bacterial load 8 hours post-infection– the time point around which mortality sets in, as described in Shit et al. (2022). In short, we extracted the whole-body homogenate of each fly individually and plated them separately on Luria-Agar plates to count the colony-forming units (CFUs) (Siva-Jothy et al. 2018). We analysed the variations in bacterial load in age groups separately. To this end, we first pseudo-log-transformed the bacterial load data and then calculated the average CFU in the three flies from each vial. We used a generalised linear mixed-effects model, with selection regime and infection dose as fixed effects and vial identity nested within the replicate populations as a random effect (Model: *LogAvgCFU ∼ Infection dose* × *Selection regime* + 1|Replicate population/Vial, *family=negative binomial*); using “glm.nb” function in *glmmTMB* package (Bates et al. 2014), followed by a posthoc test with Tukey’s adjustment to compare between regimes across different infection doses.

For the remaining 12 flies within each vial, we scored the number of dead flies every three hours for the first three days, followed by every six hours for the next two days. To analyse the survival data, we fitted the mixed effects Cox model separately to young vs old flies, using selection regime and infection dose as a fixed effect and vial identity nested within replicate populations as a random effect (Model: *Post-infection survival ∼ Infection dose* × *Selection regime +1*| *Replicate population/ Vial*); using “coxme” function in *survminer* package) (Therneau T, 2024), followed by a posthoc test with Tukey’s adjustment. We also quantified the level of susceptibility of infected flies across experimental treatments by estimating hazard ratios (HR) of deaths occurring in each of the infection treatments vs sham-infected control flies across age groups, infection doses and selection regimes (Khan et al., 2017). Hazard ratios significantly greater than one indicated a higher mortality risk in the infected groups than in sham-infected ones.

Note that our experimental design enabled us to collect paired datasets for post-infection survival estimated as hazard ratios and bacterial load for each vial across infection doses and selection regimes. We thus analysed changes in hazard ratios with variations in bacterial load by using a generalised linear model fitted separately for young and old flies, using selection regime as a fixed effect and bacterial load as a covariate. To decrease the range of distribution, we have used the log-transformed hazard ratio data and pseudo-log-transformed bacterial load data for this analysis (Model: *Log HR ∼ Log Avg CFU* × *Selection regime, family=negative binomial*). In this model outcome, a significant interaction between bacterial load and selection regime would indicate the divergence in the rate at which health declines as a function of increasing microbial load across selection regimes (i.e., a measure of infection tolerance) (Ayres and Schneider, 2012; Gupta and Vale, 2017). We used R version 4.2.2 for all the analyses.

## RESULTS

### Flies adapted to chronic malnourishment showed better post-infection survival at an early age

We found that the effects of bacterial infection on post-infection survival across selection regimes varied in a dose-dependent manner (Fig 1A, Table S1). For example, when exposed to lower infection doses, young flies from control and selected regimes showed similar susceptibility to *P. rettgeri* infection. However, at higher infection doses, flies adapted to chronic malnutrition had higher survival rates than their control counterparts. Despite the difference in post-infection survival, flies from both regimes had similar bacterial loads, indicating potential differences in infection tolerance (Fig 1C, Table S2).

**Figure 1.**
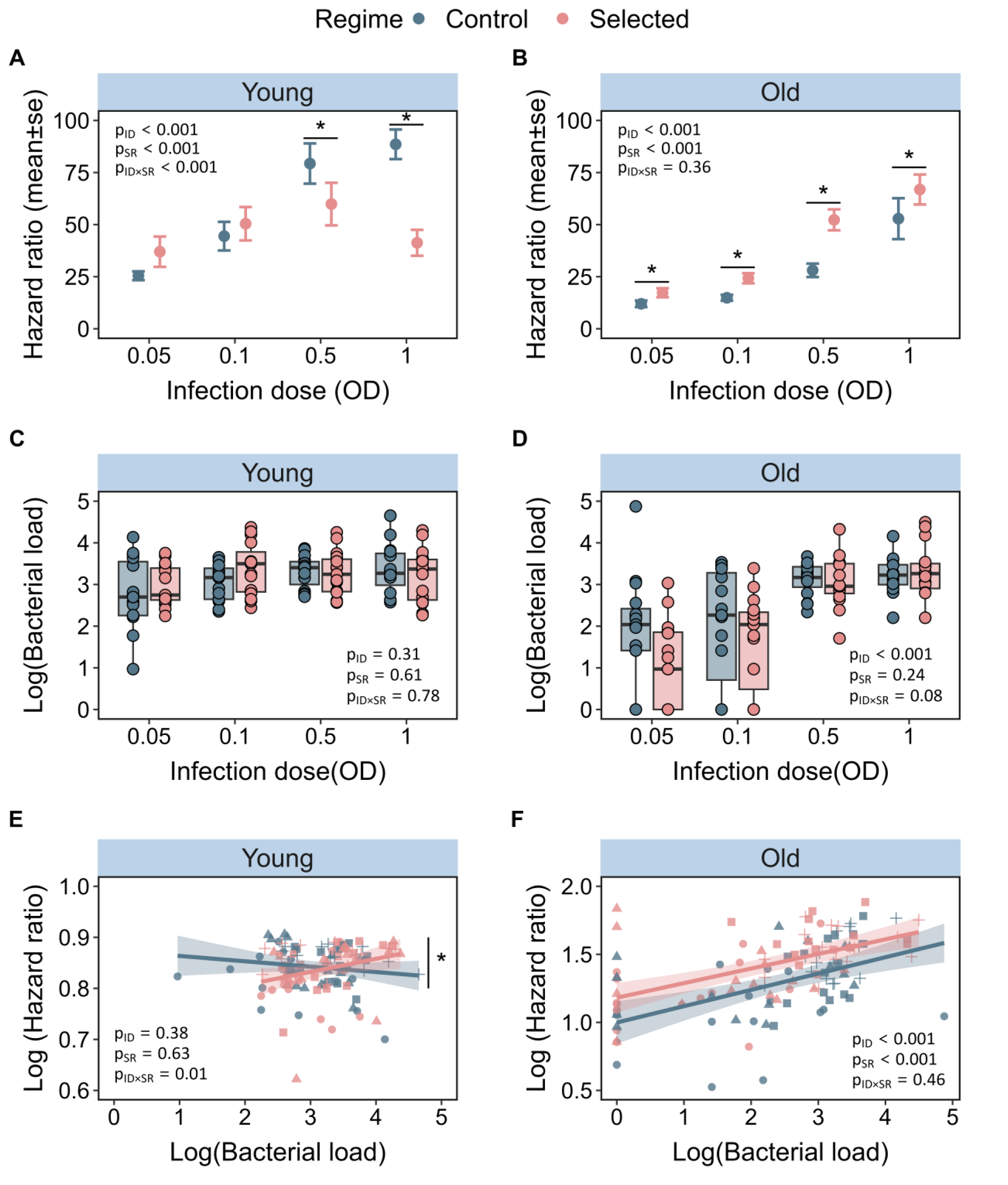
Hazard ratios for the post-infection survival response of **(A)** young and **(B)** old flies against *Providencia rettgeri* infection in control vs selected population adapted to chronic juvenile malnutrition. The estimated hazard ratios were calculated from survival curves till 120 h post-infection. Each dot represents the mean hazard ratio of five replicate vials of flies that were assayed for their survival within each experimental treatment (i.e., n= 15 females/ replicate vials/ infection dose/ age points/ selection regime/ replicate populations); Bacterial load measured for **(C)** young and **(D)** old flies across selection regimes at 8 h post-infection. Each dot represents a log-transformed Mean±SE of three individual flies sampled from each of the 5 replicated vials across selection regimes and replicate populations (n=3 females homogenised individually / replicate vials/ infection dose/ age points/ selection regime/ replicate populations); Changes of post-infection survival estimated as hazard ratios vs the corresponding bacterial load (i.e., variation in post-infection health with changes in pathogen burden; or infection tolerance) in young **(E)** and old **(F)** flies pooled across selection regimes for each replicate population. Each shape (circle=0.05 OD, triangle=0.1 OD, square =0.5 OD, plus =1 OD) represents the log-transformed hazard ratio and the correlated estimate of pseudo-log-transformed bacterial load derived from each replicate vial. In panel A-E, statistically significant differences between regimes have been indicated with an asterisk (*) (i.e., p < 0.05). Infection Dose is designated by ID and Selection Regime is designated by SR.

This is further supported by analyzing the divergent slopes of post-infection survival-vs-bacterial load data across selection regimes (characterized by a significant interaction between bacterial load and selection regime), which revealed that control flies had a higher mortality risk (i.e., higher hazard ratio of infection vs. sham-infection treatment) relative to selected flies with increasing bacterial burden inside their bodies (Fig 1E, Table S3).

### Impacts on post-infection survival were reversed with ageing

In contrast to the pattern observed at a young age, selected flies showed a significantly higher susceptibility to infection as they aged, compared to their control counterparts (1B, Table S1), in all the tested infection doses regardless of the infection doses. However, we observed no differences in their bacterial load (1D, Table S2). Also, the slope of post-infection survival-vs-bacterial load data was not detectably different across selection regimes (Table S3), suggesting that the risk of post-infection mortality did not change with variations in within-host pathogen burden across selection regimes (1F), as selected flies were consistently more susceptible than control flies.

## DISCUSSION

In this work, we investigated how long-term adaptation to early-life chronic malnutrition influences the ability to combat pathogen exposure in adulthood. Our results indicated an age-dependent effect: an early-life survival advantage that diminishes with age, increasing late-life infection susceptibility. We discovered that flies whose ancestors evolved under chronic juvenile malnutrition displayed increased survival against systemic infection by *P. rettgeri* during early adulthood. Although the overall bacterial load did not appear to change across infection doses, the evolved populations showed a relatively lower mortality increase with increasing pathogen burden, indicating higher infection tolerance (Ayres & Schneider, 2012). A prolonged evolutionary history of adapting to dietary stress may prepare organisms to develop a general stress response in evolved flies (Sinclair et al., 2013), which could help them better tolerate infection stress. Alternatively, the observed pattern could be due to altered evolutionary trade-offs between immunity and other life-history traits because of limited resource availability (Stearns, 1989). For example, hosts, when faced with pathogen risk, can reallocate their resources from traits such as reproduction to immunity and post-infection survival. Indeed, while a previous study by Kloss et al. (2009) found no adverse effects of adaptation to dietary stress on traits such as egg to adult viability or growth rate when assessed on standard food conditions in these populations, there was a notable decrease in fecundity at an early reproductive age compared to controls. Therefore, it is possible that over many generations, these flies, while evolving strategies to obtain nourishment from a poor food patch, could allocate resources to other fitness-related traits such as immune defense in the event of a pathogen attack.

Another striking aspect of our result was that the early-life survival advantage of selected flies against pathogenic *P. rettgeri* infection was reversed as the flies aged. Although flies from both selection and control regimes were equally effective at resisting and tolerating infections, we found that selected flies were generally more susceptible to infections with ageing, regardless of the initial infection dose. Despite carrying similar pathogen loads as control populations, higher fitness costs in our experimentally evolved older flies may indicate relatively higher immunopathological damage associated with their immune activation (Khan et al. 2017). Such exacerbated late-life infection costs in older flies might be attributed to the selection pressure imposed during poor nutrition in the juvenile stage that exclusively targeted early-life fitness traits (Stearns, 1989), but not late-life performance. Early-life beneficial alleles here may have traded off with late-life fitness where the selection strength was too weak to maintain the adequate level of physiological homeostasis in aged flies (Rauser et al., 2006), thereby leading to detrimental pleiotropic effects on late-life somatic maintenance during infection.

We also note that our results differ from a previous study by Vijendraverma et al. (2015) on the same malnutrition-adapted fruit fly populations, which showed that evolved flies succumbed more to *Pseudomonas entomophila* infection than controls. However, our study and that of Vijendraverma et al. (2015) differ not only in the identity of the pathogen but more importantly the route of infection and the mechanism of virulence manifestation. In the former study, flies were ingested with *P. entomophila* cells, and thus the initial immune response depended on immune activation in the gut. Toxins produced by the pathogen and possibly over-activation of ROS-induced damage made the gut wall leaky. As a consequence, the bacteria (including members of microbiota) leaked into hemolymph by breaching the gut barrier and evaded the activation of gut immunity (e.g., no changes in antimicrobial peptides across selection regimes), thereby causing septic shock (Vijendraverma et al. 2015). In contrast, flies in this experiment were systemically infected using septic injury, where *P. rettgeri* cells were introduced directly into the hemolymph, which is likely to cause rapid activation of antimicrobial peptides (Shit et al. 2022) and cause fatal septicemia (Msaad Guerfali et al., 2018). The divergent levels of immune responses deployed may thus explain the observed difference between the two studies. However, more experiments are needed to test these possibilities across pathogens and their mode of infection.

To conclude, we highlight that animals, including human ancestors (Prentice, 2005), are exposed to natural selection driven by nutrient shortages in many parts of the world. Our results provide novel insights into understanding the combined evolutionary consequences on the immune system and the ability to withstand pathogens. They also provide empirical evidence of how evolutionary ancestry and juvenile nutrition can regulate vulnerability to adult diseases with ageing, which may instigate more gerontological research to reveal the underlying genetics and metabolic changes.

## Supporting information

Sarkar et al 24_Malnutriton_Supplementory Information

## Data accessibility

Raw data will be available in the Dryad Digital Repository.

## Authors’ Contribution

Conceptualisation: IK, SS

Design of the experiment: IK, SS, BS

Funding acquisition: IK

Investigation: SS, BS, JB, SD

Supervision: IK

Data curation and analysis: SS

Writing – original draft: SS, IK

Writing – review & editing: IK, SS, TJK

## Acknowledgements

We thank Srijan Seal, Selah Makinishi and Hitee Bhardwaj for their important feedback on the manuscript. We are grateful to Tadeusz J. Kawecki for generously providing us with the fly lines.

