## Supplementary material for "Evolutionary history with chronic malnutrition enhances pathogen susceptibility at older ages": Sarkar et al 24_Malnutriton_Supplementory Information

**SUPPLEMENTARY INFORMATION**

**TABLES**

**Table S1**. Summary of a mixed effect Cox model, using selection regime and infection dose as fixed effects and vial identity nested within replicate populations as a random effect (Model: *Post-infection survival ~ Infection dose × Selection regime + 1| Replicate population/ Vial*). Statistically significant p-values are highlighted in bold.

| | | **Young** | | | | **Old** | | | |
|---|---|---|---|---|---|---|---|---|---|
| | ***Tested effect*** | *Chisq* | *Df* | *p value* | | *Chisq* | *Df* | *p value* | |
| *Full model* | *Infection dose (ID)* | 395.456 | 4 | **<2.2e-16*** ** | | 405.748 | 4 | **<2.2e-16*** ** | |
| | *Selection Regime (SR)* | 15.640 | 1 | **7.662e-05*** ** | | 65.7112 | 1 | **5.221e16*** ** | |
| | *ID ⨯ SR* | 31.344 | 4 | **2.605e-06*** ** | | 4.3667 | 4 | 0.3586 | |
| *Post hoc to compare regime across doses* | *ID* | *Estimate* | *SE* | *Z ratio* | *p-value* | *Estimate* | *SE* | *Z ratio* | *p-value* |
| | *0.05 OD* | -0.308 | 0.111 | -2.788 | 0.14 | -0.441 | 0.131 | -3.360 | **0.027** |
| | *0.1 OD* | 0.242 | 0.106 | 2.295 | 0.392 | -0.455 | 0.122 | -3.731 | **0.007** |
| | *0.5 OD* | 0.447 | 0.103 | 4.338 | **0.001** | -0.640 | 0.113 | -5.677 | **<0.001** |
| | *1 OD* | 0.409 | 0.104 | 3.948 | **0.003** | -0.405 | 0.109 | -3.725 | **0.008** |

**Table S2**. Summary of a generalised linear mixed-effects model, with selection regime and infection dose as fixed effects and vial identity nested within the replicate populations as a random effect (Model: Log Avg *CFU~ Infection dose × Selection regime + 1|Replicate population/Vial, family=negative binomial*). Statistically significant p-values have been highlighted in bold.

| | | **Young** | | | | **Old** | | | |
|---|---|---|---|---|---|---|---|---|---|
| | ***Tested effects*** | *Chisq* | *Df* | *p-value* | | *Chisq* | *Df* | *p-value* | |
| *Full model* | *Infection dose (ID)* | 3.548 | 3 | 0.314 | | 28.508 | 3 | **2.842e-06*** ** | |
| | *Selection Regime (SR)* | 0.258 | 1 | 0.612 | | 1.361 | 1 | 0.24331 | |
| | *ID ⨯ SR* | 1.074 | 3 | 0.783 | | 6.596 | 3 | 0.08595 | |
| *Post hoc to compare regime across doses* | *ID* | *Estimate* | *SE* | *Z ratio* | *p-value* | *estimate* | *SE* | *Z ratio* | *p-value* |
| | *0.05 OD* | -0.075 | 0.103 | -0.729 | 0.996 | 0.652 | 0.237 | 2.746 | 0.109 |
| | *0.1 OD* | -0.108 | 0.143 | -0.759 | 0.9950 | 0.201 | 0.258 | 0.776 | 0.994 |
| | *0.5 OD* | 0.014 | 0.152 | 0.092 | 1.000 | 0.008 | 0.199 | 0.039 | 1.000 |
| | *1 OD* | 0.043 | 0.149 | 0.288 | 1.000 | -0.021 | 0.194 | -0.107 | 1.000 |

**Table S3**. Summary of the generalised linear model fitted separately for young and old flies, using log-transformed hazard function (HR) as a response variable, pseudo-log-transformed bacterial load (CFU) as a covariate and selection regime (SR) as a fixed effect (model: *LogHR~ Log Avg CFU* × *Selection regime*, *family=negative binomial*). A significant interaction between bacterial load and selection regime indicates variation in infection tolerance. Statistically significant P-values have been highlighted in bold.

| | ***Young*** | | | ***Old*** | | |
|---|---|---|---|---|---|---|
| ***Tested effect*** | *Chisq* | *Df* | *p-value* | *Chisq* | *Df* | *p-value* |
| *LogAvgCFU* | 0.7710 | 1 | 0.37990 | 42.836 | 1 | **5.951e-11** *** |
| *Selection regime (SR)* | 0.2294 | 1 | 0.63194 | 12.074 | 1 | **0.0005112** *** |
| *LogAvgCFU* × *SR* | 6.3165 | 1 | **0.01196*** | 0.545 | 1 | 0.4602286 |

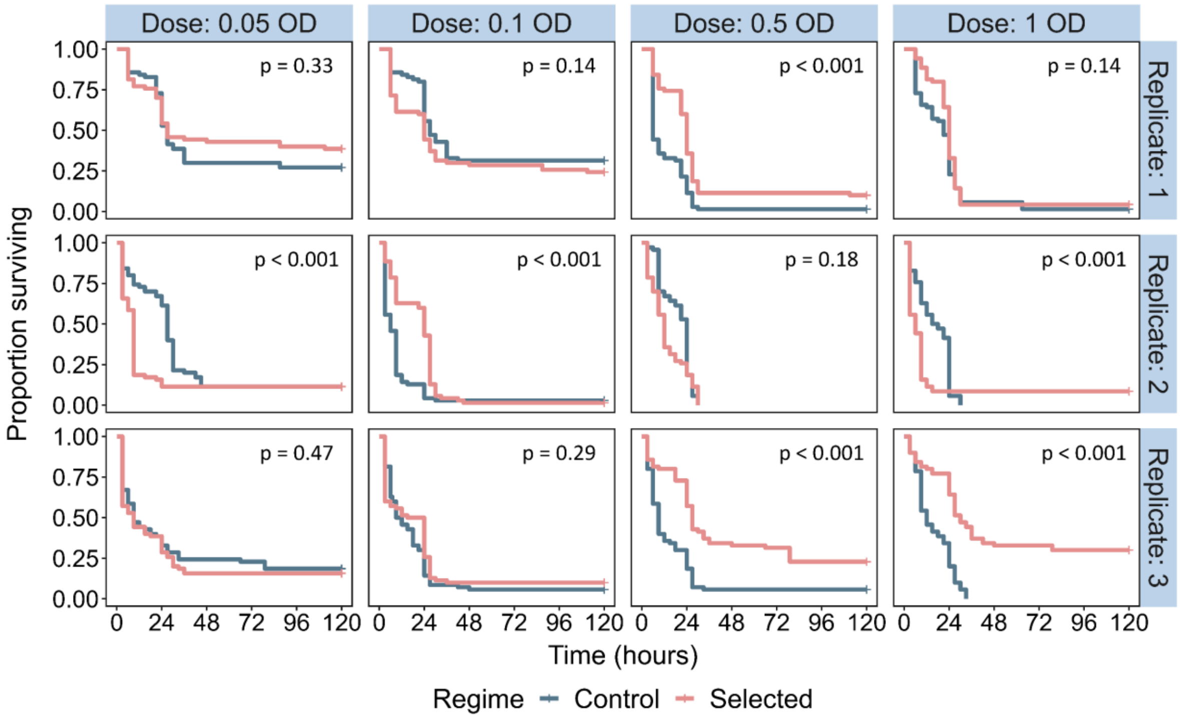


**Figure S1.** Replicate population data for post-infection survival of selected flies and their control 120 hours post *P. rettgeri* infection at a young age.

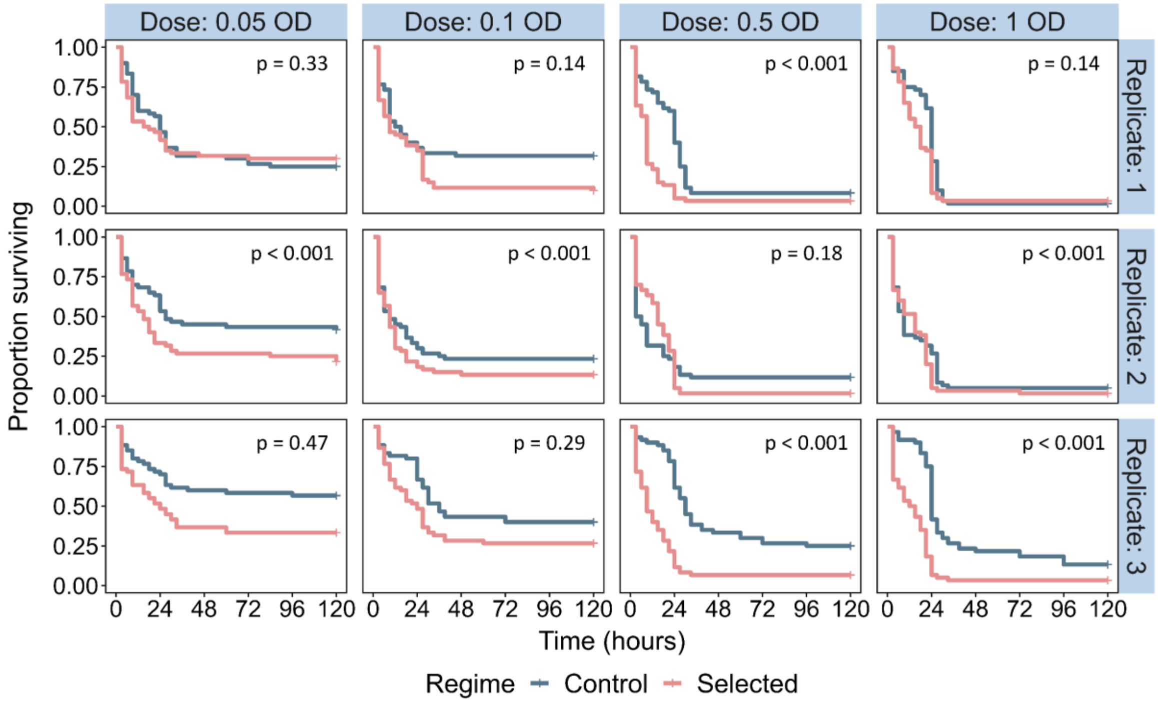


**Figure S2.** Replicate population data for post-infection survival of selected flies and their control 120 hours post *P. rettgeri* infection at old age.

**Table S4**. The effect of infection dose and selection regime on survival of individual replicate fly population 120 h post-infection with *Providencia rettgeri* at a young age. Values in bold are statistically significant.

| | | | | | | | |
|---|---|---|---|---|---|---|---|
| | | | *Tested effect* | | *Chisq* | *Df* | *p-value* |
| ***Young*** | *Replicate set 1* | *Full model* | *Dose* | | 199.703 | 4 | **< 2.2e-16 ***** |
| | | | *Regime* | | 10.639 | 1 | **0.0011072**** |
| | | | *Dose* ╳ *Regime* | | 21.895 | 4 | **0.0002103***** |
| | | Dose specific comparisons across regimes | *Dose* | *loglik* | *Chisq* | *Df* | *p-value* |
| | | | *0.05 OD* | -421.78 | 0.9658 | 1 | 0.3257 |
| | | | *0.1 OD* | -447.49 | 2.1946 | 1 | 0.1385 |
| | | | *0.5 OD* | -532.66 | 23.905 | 1 | **1.012e-06 ***** |
| | | | *1 OD* | -550.93 | 2.2173 | 1 | 0.1365 |
| | *Replicate set 2* | *Full model* | *Dose* | | 168.964 | 4 | **<2.2e-16***** |
| | | | *Regime* | | 1.237 | 1 | **0.266** |
| | | | *Dose* ╳ *Regime* | | 58.326 | 4 | **6.518e-12***** |
| | | Dose specific comparisons across regimes | *Dose* | *loglik* | *Chisq* | *Df* | *p-value* |
| | | | *0.05 OD* | -517.48 | 14.129 | 1 | **0.0001707 ***** |
| | | | *0.1 OD* | -541.47 | 23.907 | 1 | **1.011e-06 ***** |
| | | | *0.5 OD* | -554.32 | 1.793 | 1 | 0.1806 |
| | | | *1 OD* | -543.72 | 9.839 | 1 | **0.001708 **** |
| | *Replicate set 3* | *Full model* | *Dose* | | 127.469 | 4 | **< 2.2e-16 ***** |
| | | | *Regime* | | 25.471 | 1 | **4.491e-07 ***** |
| | | | *Dose* ╳ *Regime* | | 35.697 | 4 | **3.341e-06 ***** |
| | | Dose specific comparisons across regimes | *Dose* | *loglik* | *Chisq* | *Df* | *p-value* |
| | | | *0.05 OD* | -500.17 | 0.5282 | 1 | 0.4674 |
| | | | *0.1 OD* | -537.16 | 1.1202 | 1 | 0.2899 |
| | | | *0.5 OD* | -500.24 | 25.291 | 1 | **4.931e-07 ***** |
| | | | *1 OD* | -487.50 | 44.678 | 1 | **2.322e-11 ***** |

**Table S5**. The effect of infection dose and selection regime on survival of individual replicate fly population 120 post-infection with *Providencia rettgeri* at an old age. Values in bold are statistically significant.

| | | | *Tested effect* | | *Chisq* | *Df* | *p-value* |
|---|---|---|---|---|---|---|---|
| ***Old*** | *Replicate set 1* | *Full model* | *Dose* | | 134.354 | 4 | **< 2.2e-16 ***** |
| | | | *Regime* | | 18.979 | 1 | **1.322e-05***** |
| | | | *Dose ╳ Regime* | | 10.330 | 4 | **0.03522*** |
| | | Dose specific comparisons across regimes | *Dose* | *loglik* | *Chisq* | *Df* | *p-value* |
| | | | *0.05 OD* | -0.372.72 | 0.0743 | 1 | 0.7852 |
| | | | *0.1 OD* | -0.397.42 | 4.779 | 1 | **0.02881 *** |
| | | | *0.5 OD* | -438.24 | 22.092 | 1 | **2.599e-06 ***** |
| | | | *1 OD* | -453.00 | 6.0421 | 1 | **0.01397 *** |
| | *Replicate set 2* | *Full model* | *Dose* | | 137.049 | 4 | **<2.2e-16***** |
| | | | *Regime* | | 5.081 | 1 | **0.02419*** |
| | | | *Dose ╳ Regime* | | 5.048 | 4 | 0.28242 |
| | | Dose specific comparisons across regimes | *Dose* | *loglik* | *Chisq* | *Df* | *p-value* |
| | | | *0.05 OD* | -351.72 | 6.2551 | 1 | **0.01238 *** |
| | | | *0.1 OD* | -408.47 | 1.7326 | 1 | 0.1881 |
| | | | *0.5 OD* | -447.21 | 0 | 1 | 0.996 |
| | | | *1 OD* | -453.79 | 1.6845 | 1 | 0.1943 |
| | *Replicate set 3* | *Full model* | *Dose* | | 143.624 | 4 | **< 2.2e-16 ***** |
| | | | *Regime* | | 60.324 | 1 | **8.045e-15 ***** |
| | | | *Dose ╳ Regime* | | 6.552 | 4 | 0.1616 |
| | | Dose specific comparisons across regimes | *Dose* | *loglik* | *Chisq* | *Df* | *p-value* |
| | | | *0.05 OD* | -293.49 | 7.1567 | 1 | **0.007469 **** |
| | | | *0.1 OD* | -345.34 | 4.3112 | 1 | **0.03786 *** |
| | | | *0.5 OD* | -402.18 | 32.582 | 1 | **1.143e-08 ***** |
| | | | *1 OD* | -426.61 | 32.19 | 1 | **1.398e-08 ***** |
